# Developmental shift in β-catenin localization between nuclear and junctional pools during vertebrate nephron development

**DOI:** 10.64898/2026.07.23.740327

**Authors:** Adrian Romero, Amaya C. Moss, Brandy L. Walker, Ulrich Rothbauer, Rachel K. Miller

## Abstract

Wnt/β-catenin signaling is a critical pathway that regulates nephron progenitor renewal versus differentiation as well as nephron patterning. In addition to its role as a transcription co-factor, β-catenin also functions as a structural component of adherens junctions, where it interacts with cadherins to link cell-cell contacts to the cytoskeleton. However, the relationship between the nuclear and junctional localization of β-catenin during vertebrate nephron development remains poorly understood. To define how endogenous β-catenin localization changes during nephrogenesis, we optimized an accelerated-turnover β-catenin chromobody for live imaging in *Xenopus* embryos. Using *in vivo* imaging of *Xenopus* pronephric development, we visualized endogenous β-catenin within the nuclear, cytoplasmic, and junctional compartments. Across successive developmental stages, β-catenin became progressively enriched at epithelial junctions during nephron maturation while remaining abundant within nuclear and cytoplasmic compartments. Quantitative analyses indicate that epithelial maturation is accompanied by coordinated expansion and partitioning of multiple intracellular β-catenin pools rather than a simple redistribution from nuclear to junctional compartments.

## 1. Introduction

Nephrons are the structural and functional units of vertebrate kidneys, forming a highly segmented epithelial network that mediates precise fluid filtration and homeostasis. These units develop through coordinated interactions between specific cells called nephron progenitors, which differentiate into epithelial lineages through mesenchymal-to-epithelial transitions (1–3). Therefore, understanding how those nephron segments are first specified and subsequently organized during kidney development is essential to define the mechanisms that establish a functional kidney (4–7).

During development, the kidney arises from the intermediate mesoderm. In amniotes, kidney formation occurs in three successive phases. The development of the embryonic pronephros and mesonephros, which are transient structures, precedes the formation of the metanephros, the adult kidney. In anamniotes, the embryonic pronephros is functional and is the foundation for the development of the adult mesonephros. The nephrons within the forms share conserved cellular and molecular features with mammalian nephrons, making them useful models for studying vertebrate nephrogenesis (3, 4). In the *Xenopus* model, the pronephric kidney consists of a single nephron on each side of the embryo. It is segmented into proximal and distal tubules, a connecting tubule, and the nephric duct. Several genes involved in kidney development in amphibians are well conserved, supporting their use for functional genetic studies and disease modeling (8, 9). In addition, the large external development of the embryo makes it easily accessible for targeted microinjection and suitable for live imaging, making *Xenopus* particularly useful for studying protein localization and cellular behaviors during organogenesis (4, 10, 11).

Canonical Wnt/β-catenin signaling plays a central role in nephron specification and epithelial differentiation. In the absence of Wnt ligands, cytoplasmic β-catenin is continuously phosphorylated by the destruction complex, composed of Axin, APC, CK1, and GSK3β, targeting it for ubiquitin-mediated degradation. Activation of Wnt signaling inhibits this destruction complex, allowing β-catenin to accumulate in the cytoplasm and translocate to the nucleus, where it associates with TCF/LEF transcription factors to regulate target gene expression (12–16). In addition to its transcriptional role, β-catenin is an essential structural component of adherens junctions. At the plasma membrane, β-catenin binds classical cadherins and links epithelial cell-cell junctions to the actin cytoskeleton through α-catenin (17–19). Consequently, β-catenin continuously distributes among nuclear, cytoplasmic, and junctional pools depending on developmental stage and cellular context. Because these pools reflect distinct biological functions, characterization of β-catenin subcellular localization provides information that cannot be obtained solely from gene expression or total protein abundance measurements.

The role of canonical Wnt/β-catenin signaling during kidney development is well understood. In the mammalian metanephros, Wnt9b initiates nephron induction by promoting the mesenchymal-to-epithelial transition of nephron progenitors, whereas Wnt4 reinforces epithelial differentiation and nephron formation (20–23). In the pronephric *Xenopus* kidney, Wnt4 is required for tubulogenesis and nephron patterning, and disruption of canonical Wnt signaling results in severe defects in kidney epithelial development (24–26). Additional studies have identified several Wnt ligands expressed within the developing pronephros, including *wnt4*, *wnt9a*, and *wnt11*, indicating that canonical Wnt signaling is spatially regulated during nephrogenesis (2, 27–30). However, considerably less is known about how endogenous β-catenin protein is dynamically distributed among its different subcellular compartments during nephrogenesis.

Most previous studies have relied on pathway perturbations, transcriptional reporters, or fixed-tissue immunostaining, approaches that provide limited information regarding the dynamic partitioning of β-catenin among its signaling and structural pools in living embryos. Chromobody technology provides an opportunity to visualize endogenous proteins *in vivo* (31, 32). Chromobodies consist of fluorescently labeled single-domain antibody fragments derived from camelid heavy-chain antibodies that bind intracellular target proteins without requiring fixation (32). The β-catenin chromobody BC1-CB recognizes endogenous β-catenin and has previously been used to monitor β-catenin localization and nuclear translocation in cultured cells (31, 32). However, its application to vertebrate kidney development has not been investigated.

We hypothesized that endogenous β-catenin undergoes developmentally regulated partitioning between transcriptional and adhesive pools during nephrogenesis. To test this hypothesis, we adapted an accelerated-turnover β-catenin chromobody for live imaging of *Xenopus* kidney development. We show that β-catenin becomes progressively enriched at epithelial junctions during tubule maturation, while remaining readily detectable in both nuclear and cytoplasmic compartments. Quantitative analyses indicate that nephrogenesis is accompanied by developmental partitioning of multiple intracellular β-catenin pools rather than a simple transition from nuclear to junctional localization.

## 2. Methods

### 2.1 Xenopus laevis embryo collection and culture

*Xenopus laevis* frogs were purchased from NXR and *Xenopus* 1 and maintained according to standard animal husbandry protocols. All animal work was conducted in accordance with IACUC-approved protocol (AWC-25-0074) from the University of Texas Health Science Center at Houston. Female frogs were induced with human chorionic gonadotropin (hCG), and eggs were collected 14–16 hours later by manual squeezing. Eggs were fertilized using macerated male testis in 1x MMR. After fertilization, embryos were maintained in 1/3x MMR and allowed to undergo cortical rotation (30 min) before the jelly coat was removed using 2% cysteine solution for 5 min. Embryos were staged according to Nieuwkoop and Faber(33) and sorted for viability before microinjection.

### 2.2 mRNA synthesis, and microinjection

To enable expression in *Xenopus* embryos, previously published β-catenin chromobody constructs (34, 35) were subcloned into the pCS2+ backbone. The resulting plasmids were linearized with *NotI*. Capped mRNA was synthesized using the SP6 mMessage kit and purified by phenol/chloroform extraction and precipitation. The β-catenin chromobody mRNA and the membrane tracer mRNA were injected at 300 pg by embryo respectively. Embryos were injected at the 8-cell stage into the V2 blastomere to target kidney fated cells. The injection volume was kept consistent at 10 nL. Each experiment used approximately 50 embryos by condition. Injected embryos were sorted by fluorescence using an epifluorescence stereoscope by the membrane tracer. Embryos were analyzed at several stages to evaluate β-catenin localization during early development, epithelialization and tadpole stages when the pronephric kidney becomes functional.

### 2.3 Whole-mount immunostaining

Embryos were fixed in 4% paraformaldehyde, washed in PBS, and permeabilized in PBS-T containing 0.1% Triton. Embryos were blocked in 20% goat serum and incubated with primary antibodies overnight at 4 °C. The primary antibodies and stains used were anti-GFP chicken polyclonal to detect GFP expression (1:500; Abcam, ab13970), β-catenin mouse monoclonal antibody (1:250; Sigma-Aldrich, C7207), Lhx1 rabbit antibody (36) (1:250), and 3G8/4A6 antibodies to label proximal tubule lumens and cell membranes of the intermediate, distal, and connecting tubules respectively (1:5 and 1:30; European *Xenopus* Resource Centre). Proximal tubule lumens were also detected using FITC-conjugated lectin from *Erythrina cristagalli* (1:1000; FL-1141, Vector Laboratories). E-cadherin mouse antibody was used to label epithelial membranes in the mature pronephric kidney (BD Biosciences, 610182). DAPI was used at 1:1000 to stain nuclei. After primary antibody incubation, embryos were washed and incubated with secondary antibodies overnight at 4 °C. Goat anti-mouse, goat anti-rabbit, or goat anti-chicken secondary antibodies conjugated to Alexa Fluor 488, Alexa Fluor 555, or Alexa Fluor 647 were used at 1:500. Embryos were washed in PBS-T and PBS, dehydrated in methanol, and cleared in BABB before imaging.

### 2.5 Imaging quantification and statistical analysis

The β-catenin chromobody was used to visualize endogenous β-catenin localization *in vivo* and in fixed tissues. The chromobody recognizes the N-terminal region of β-catenin and has been reported to monitor β-catenin localization and nuclear translocation in living cells (31, 32, 34, 37). Confocal images were taken in many stages and were analyzed using Fiji/ImageJ. For each experiment, raw grayscale channels were used for quantification. When needed, z-stacks were projected using maximum-intensity projections for nuclear segmentation and sum-intensity projections for signal quantification. Kidney regions were defined manually using the corresponding kidney progenitor marker for early stages and, when available, the epithelial marker for late stages. In fixed samples, nuclear β-catenin chromobody signal was quantified using DAPI-defined nuclear regions of interest. Nuclear masks were generated from the nuclear channel and applied to the β-catenin chromobody channel to measure mean intensity and integrated density. For nuclear intensity analysis, β-catenin chromobody mean intensity was normalized to DAPI mean intensity for each nucleus and reported as nuclear β-catenin/DAPI.

For compartmentalization analysis, β-catenin chromobody integrated density was measured within DAPI-defined nuclei and compared with total β-catenin chromobody integrated density within the pronephric region. The nuclear fraction was calculated as nuclear β-catenin integrated density divided by total kidney β-catenin integrated density. The non-nuclear fraction was calculated as the remaining signal outside the nuclear compartment. Because no membrane or junctional marker was present in all developmental-stage images, the non-nuclear fraction includes cytosolic, cortical, and potential membrane-associated β-catenin signal and was not interpreted as a specific junctional compartment. For late-stage epithelial colocalization analysis, epithelial regions were identified using E-cadherin staining. β-catenin chromobody signal associated with epithelial marker-positive regions was quantified from the β-catenin channel using marker-defined regions of interest. Colocalization between β-catenin chromobody and the epithelial marker was analyzed in Fiji using Coloc 2. Pearson correlation coefficients were used to estimate global channel association, while thresholded Manders coefficients were used to estimate signal overlap. tM1 represents the fraction of thresholded β-catenin chromobody signal overlapping epithelial marker-positive regions, and tM2 represents the fraction of thresholded epithelial marker signal overlapping β-catenin chromobody.

For proximal and distal tubule analysis, 3G8 and 4A6 staining were made. DAPI-defined nuclear ROIs located within each tubule segment were used to measure nuclear β-catenin chromobody intensity. β-catenin nuclear intensity was normalized to DAPI intensity for each nucleus. Proximal and distal nuclear β-catenin/DAPI values were compared using a Wilcoxon rank-sum test. All quantitative tables exported from Fiji were analyzed in R. Graphs were generated using ggplot2. For developmental-stage compartmentalization analyses acquired under different imaging sessions and confocal settings, values were presented descriptively and no statistical comparisons were performed across stages.

## 3. Results

### 3.1 Evaluation of β-Catenin Chromobody localization in embryonic tissues

Before using the β-catenin chromobody to study pronephric kidney development, we first evaluated its localization in embryonic tissues where β-catenin activity has been previously described. We analyzed its distribution in the developing epidermis, a tissue with well-characterized β-catenin dynamics during cell fate decisions and differentiation (38). To reduce prolonged aberrant fluorescence in unbound chromobodies, a lysine residue was incorporated at the N-terminus of the mCherry- or GFP-tagged chromobody constructs to facilitate ubiquitin-mediated degradation, thereby promoting accelerated turnover (39). Blastomeres contributing to the epidermal lineage were microinjected with this accelerated-turnover β-catenin chromobody construct with a membrane tracer to track the targeted cells. Confocal imaging revealed chromobody signal across multiple subcellular compartments, including the nucleus, cytoplasm, and cell borders (Fig. S1). These observations showed that the chromobody detects β-catenin distribution without an obvious bias toward a single cellular pool.

We next evaluated β-catenin chromobody localization during early gastrulation, when dorsal β-catenin stabilization is a well-established feature of organizer formation. Gastrulation is a critical stage of embryogenesis during which the three germ layers are formed, and the basic body plan begins to take shape (40–42). During this stage, coordinated cell movements, including convergence and extension, are required for proper morphogenesis (43–45). In *Xenopus*, dorsal β-catenin stabilization contributes to early embryonic patterning and formation of the Spemann-Mangold organizer (46, 47). To evaluate chromobody localization relative to a known canonical Wnt signaling region, the dorsal side of the embryo was targeted by injection of chromobody mRNA. Embryos were allowed to develop until gastrula stages and were then analyzed for β-catenin chromobody localization. Strong β-catenin chromobody fluorescence was observed in both nuclei and cell junctions near the dorsal blastopore lip, consistent with the known role of β-catenin in dorsal organizer formation and canonical Wnt signaling (Fig. S2). Together, these validation experiments supported the use of the β-catenin chromobody as a live-imaging tool to visualize β-catenin localization *in vivo* prior to its application to the developing pronephric kidney.

### 3.2 β-Catenin Chromobody Compartmentalization Changes Between Developmental Stages

Having validated the chromobody in embryonic tissues with well-established β-catenin localization patterns, we next asked whether endogenous β-catenin partitioning changes during nephrogenesis. To evaluate β-catenin chromobody localization during pronephric kidney development, embryos injected with β-catenin-CB::EGFP were fixed at different developmental stages and analyzed by immunofluorescence. At early stages, Lhx1 was used as a nephron progenitor marker to identify the pronephric kidney region. Because Lhx1 labels nephron progenitor nuclei, it also allowed us to evaluate whether β-catenin chromobody signal was present in the nuclei of kidney cells.

Across successive developmental stages, β-catenin became progressively enriched outside the nucleus, with increasing membrane-associated localization as epithelial tubules matured, consistent with developmental partitioning of multiple intracellular β-catenin pools. At NF22, β-catenin chromobody signal was detected in nephron progenitor nuclei, with additional signal visible in the cytoplasm and in membrane-like protrusive structures (Fig. 1A). By NF26, more cells showed nuclear β-catenin signal, particularly in cells beginning to form cell contacts (Fig. 1B). These observations suggest that β-catenin is present in the early pronephric field during the transition from nephron progenitor specification toward epithelial organization.

**Figure 1.**
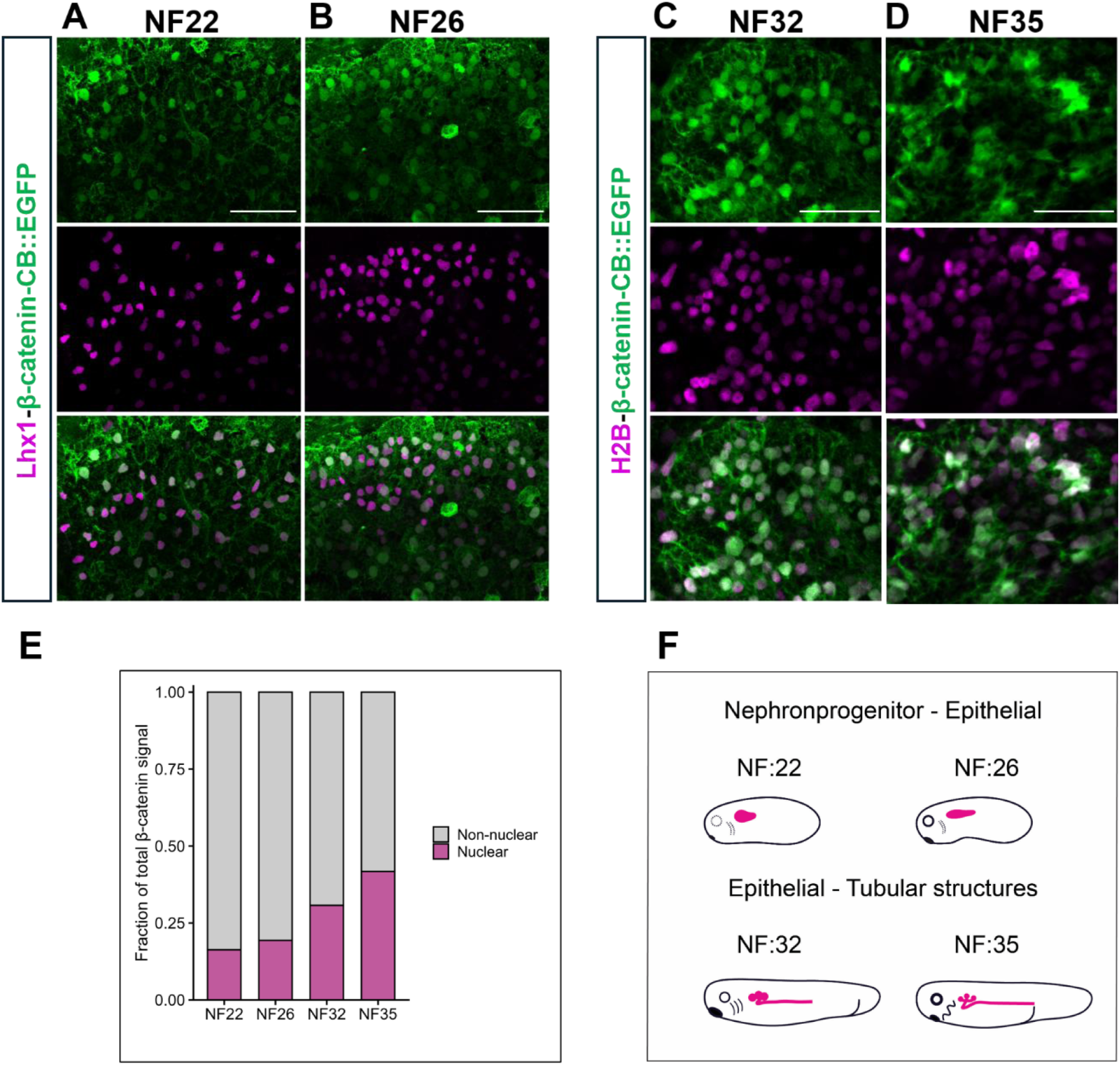
β-catenin chromobody localization in the developing *Xenopus* pronephros. Immunofluorescence analysis of embryos injected with β-catenin-CB::EGFP at various stages. A. β-catenin-CB::EGFP was detected using an anti-GFP antibody and is shown in green. Lhx1 was used as a nephron progenitor marker to identify the pronephric kidney region. Because Lhx1 labels nephron progenitor nuclei, it was also used to evaluate whether β-catenin chromobody signal was present in the nuclei of kidney cells. At NF22 β-catenin signal was detected in some nuclei. B. By NF26, more cells showed nuclear β-catenin signal, particularly in cells beginning to form border-like epithelial contacts. C. At NF32 and D. NF35, β-catenin-CB::EGFP was used to evaluate β-catenin localization during pronephric epithelialization. H2B was co-injected with the β-catenin chromobody to label nuclei. β-catenin signal was detected in nuclei during epithelialization and was also observed at cell borders as the pronephric kidney became more organized. E. Nuclear β-catenin signal was measured within DAPI-defined nuclear ROIs and expressed as a fraction of total kidney β-catenin chromobody signal. The remaining signal was classified as non-nuclear. Because no membrane marker was present throughout this experiment, the non-nuclear fraction includes cytosolic, cortical, and potentially membrane-associated signals. F. Diagrams of *Xenopus* embryos across the analyzed developmental stages, with the kidney region highlighted in pink. Scale bars: 50 µm.

For later stages, NF32 and NF35, H2B was co-injected with the β-catenin chromobody to label nuclei. At these stages, the kidney was selected based on its epithelial organization and the characteristic morphology of the developing pronephros. During pronephric epithelialization, β-catenin chromobody signal was detected in nuclei and was also observed at cell borders as the kidney became more organized, and the signal appeared to outline the simple columnar epithelial organization characteristic of epithelialized pronephric cells (Fig. 1C). At stage 35, the kidney starts to form tubular structures, and the β-catenin chromobody signal was present in all the targeted kidney cells and in a potentially membrane-associated signal (Fig. 1D).

To quantify this localization pattern, the area of the nuclear β-catenin signal was measured within DAPI-defined kidney regions (not shown) and expressed as a fraction of the total β-catenin chromobody area. The remaining signal was classified as non-nuclear. Because no membrane marker was present throughout this experiment, the non-nuclear fraction includes cytosolic, cortical, and potentially membrane-associated β-catenin signal. This analysis demonstrated that nuclear β-catenin remained a substantial component of the total β-catenin pool throughout pronephric development, while the non-nuclear pool expanded as epithelial tubules formed (Fig. 1E). To contextualize the specific developmental stages analyzed in this study, diagrams of *Xenopus* embryos highlighting the renal region (pink) are provided in Figure 1F. Together, these findings indicate that epithelial maturation is accompanied by increasing compartmentalization of β-catenin rather than replacement of one pool by another.

### 3.3 β-Catenin localization in the Xenopus Pronephric Kidney during Development In Vivo

We next asked whether the developmental expansion and dynamic partitioning of β-catenin among intracellular compartments could be visualized directly in living embryos. To determine whether the β-catenin chromobody localizes in the nephron in formation, we used β-catenin chromobody live imaging in the *pax8*-GFP transgenic frog [*Xla.Tg(Xtr.pax8:GFP)Ogino*] (48) to evaluate whether targeted kidney cells expressing β-catenin during *Xenopus* kidney development *in vivo*. These experiments were performed by injecting β-catenin chromobody mRNA into the V2 blastomere at the 8-cell stage. We performed time-lapse imaging during NF30 development, when the pronephros is starting its transition toward epithelial tubule structures. We selected this stage because the *pax8*-GFP reporter is easily visible in kidney structures and accessible through the skin. Our aim was to evaluate chromobody localization in the kidney and determine the subcellular localization of the β-catenin signal during tubule development.

At this stage, the β-catenin chromobody signal was observed in the kidney and surrounding tissue (Fig. 2A). A short time-lapse also revealed β-catenin localization dynamics throughout the pronephros during development. During the 90-minute time-lapse, some cells already showed nuclear β-catenin signal, and over time, the chromobody signal further accumulated in nuclei (Fig. 2B) or in structures such as forming cell contacts (Fig. 2C). These observations demonstrate that the chromobody enables visualization of dynamic β-catenin partitioning *in vivo* during nephron morphogenesis.

**Figure 2.**
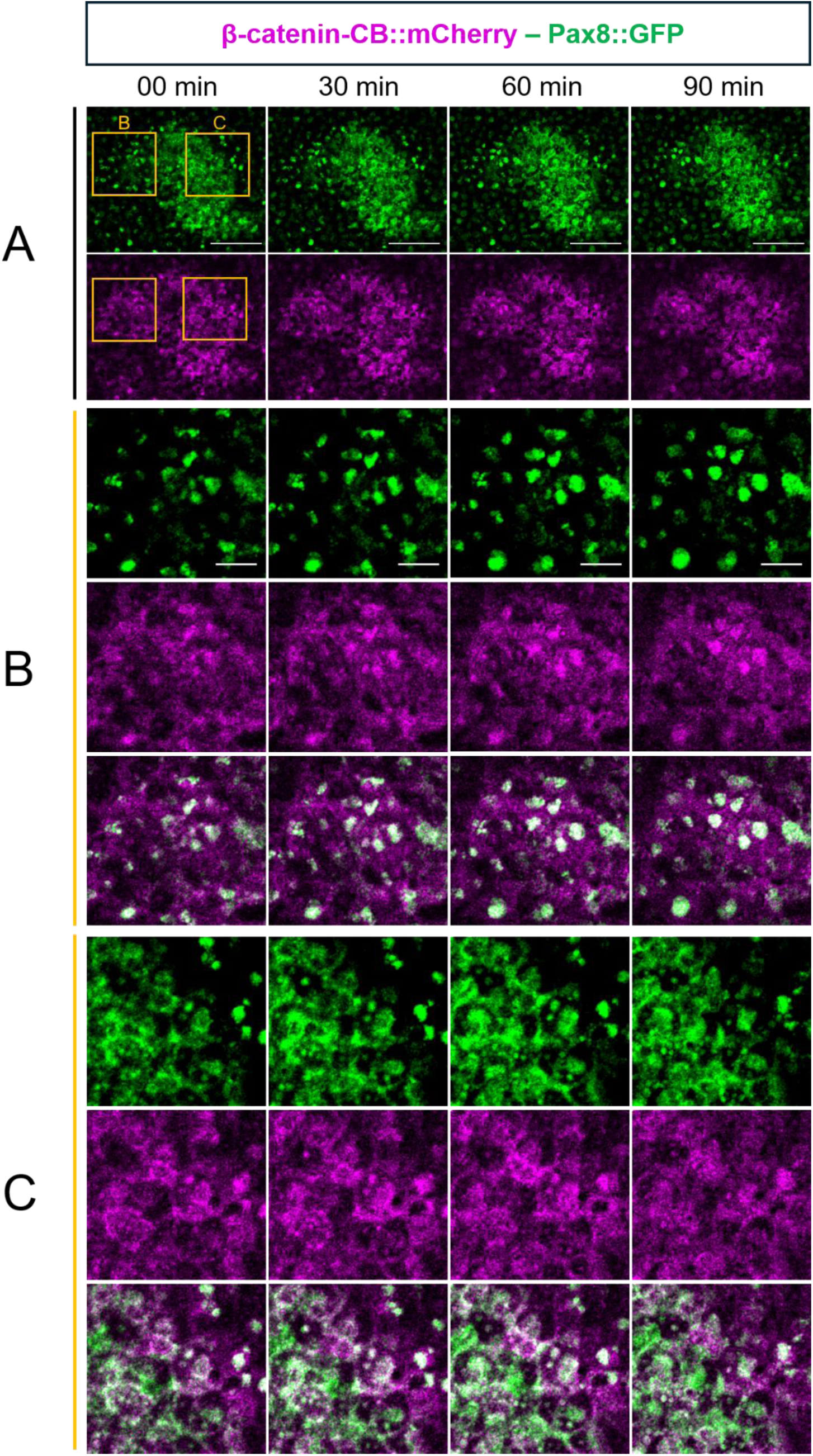
Live imaging of β-catenin chromobody localization during *Xenopus* pronephric kidney development. Time-lapse imaging shows pronephric kidney development at NF30, when kidney tubules are forming. The kidney is shown in green using a transgenic *pax8*-GFP reporter. Embryos were injected with mRNA encoding β-catenin-CB::mCherry, shown in magenta. In panel A, at 0 min, the *pax8*-GFP reporter marks the pronephric kidney, while β-catenin-CB::mCherry is detected around and inside the pronephric structure. After 90 min of time-lapse imaging, the kidney structure appears more compact, possibly due to contraction or remodeling of the developing pronephros. Two zoomed-in regions are shown in panels B and C. In A, a group of cells becomes more compact after 90 min, with circular structures resembling nuclei. The merge shows overlap between the *pax8*-GFP kidney reporter and β-catenin-CB::mCherry signal, indicating that the kidney structure was targeted. In C, it shows β-catenin chromobody dynamics over time in structures that appear cytosolic and border-like. Scale bar in panel A: 100 µm. Scale bar in panel B: 25 µm.

### 3.4 β-Catenin Chromobody Localizes to E-Cadherin-Positive Epithelial Structures in the Mature Pronephric Kidney

We next investigated later stages to understand β-catenin chromobody localization after pronephric epithelialization. These experiments were performed by injecting β-catenin-CB::EGFP into the V2 blastomere at the 8-cell stage. At NF40, E-cadherin was used to label epithelial membranes and delineate the mature pronephric kidney. At this stage, β-catenin chromobody signal was detected in multiple compartments of the pronephric tubule, including DAPI-defined nuclei, cytosolic structures, and E-cadherin-positive epithelial regions (Fig. 3A–D). To evaluate this localization more directly, we performed compartmental analysis of the β-catenin chromobody signal within the kidney field. This analysis showed that a fraction of the total kidney chromobody signal was nuclear, another fraction was associated with E-cadherin-positive regions, and the remaining signal was present in other intracellular regions within the epithelial kidney field (Fig. 3E). These observations indicate that β-catenin chromobody is not restricted to one cellular compartment at NF40, but remains distributed across nuclear, cytosolic, and epithelial-associated regions.

**Figure 3.**
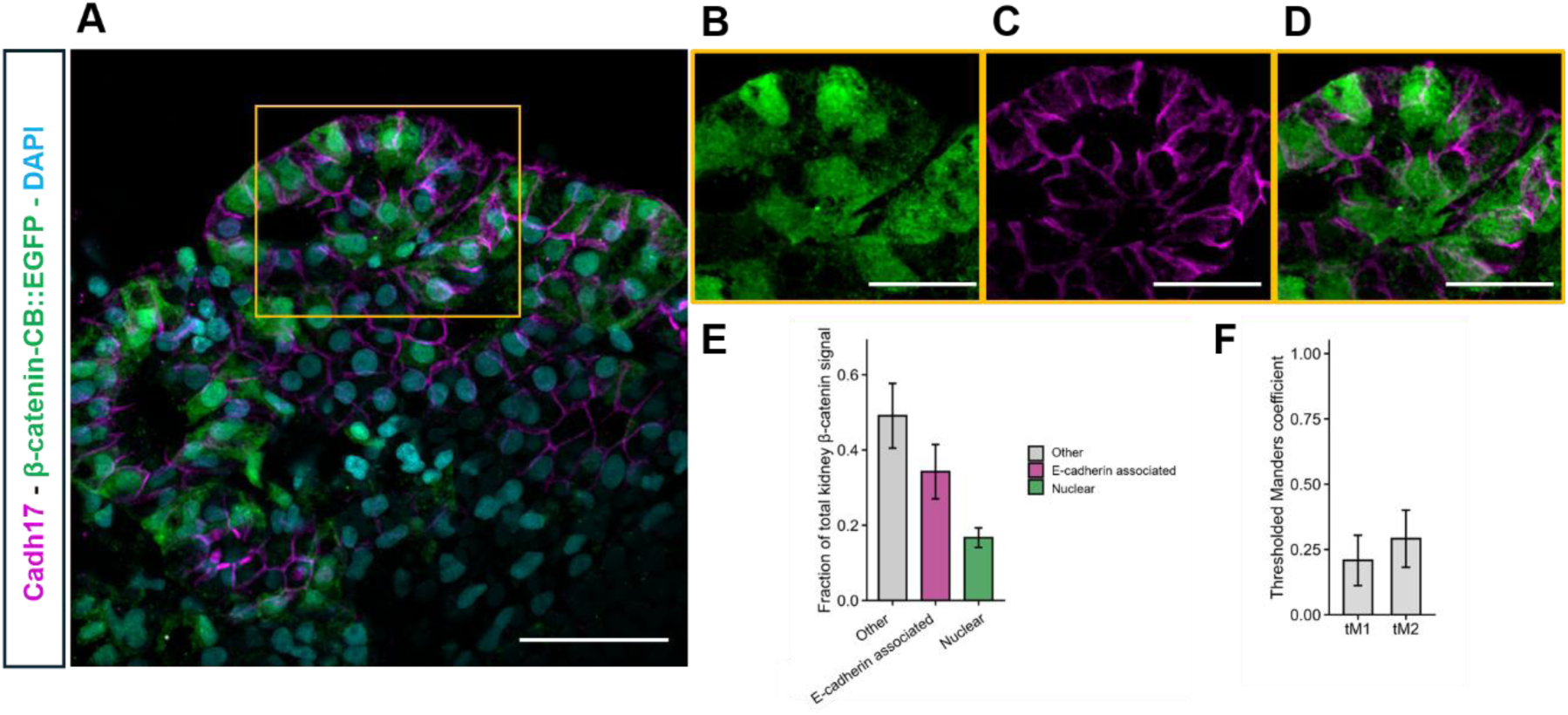
β-catenin chromobody localization in E-cadherin-positive epithelial structures of the mature pronephric kidney. At stage NF40, E-cadherin was used to label epithelial membranes and delineate the mature pronephric kidney. β-catenin chromobody signal was detected in multiple cellular compartments, including nuclei, epithelial cell borders, and cytosolic regions. Panels B–D show magnified regions highlighting β-catenin chromobody localization within E-cadherin-positive epithelial structures. In panel E, a compartmentalization analysis shows the fraction of total kidney β-catenin chromobody signal assigned to nuclei, E-cadherin-associated regions, and other intracellular regions within the epithelial kidney field. In panel F, thresholded Manders coefficients were used to estimate overlap between β-catenin chromobody and E-cadherin. tM1 indicates the fraction of β-catenin chromobody signal overlapping E-cadherin-positive regions, while tM2 indicates the fraction of E-cadherin signal overlapping β-catenin chromobody. Values are shown as mean ± SD. Scale bar in panel A: 50 µm and B-D: 25 µm.

We then quantified the overlap between β-catenin chromobody and E-cadherin in background-corrected NF40 kidney images using Coloc 2 (Fiji plugin). Pearson correlation was used to assess global channel association, while thresholded Manders coefficients were used to estimate signal overlap. Across four imaging embryos, β-catenin chromobody showed a modest-to-moderate positive association with E-cadherin. Pearson correlation values ranged from 0.20 to 0.52, while thresholded Manders coefficients showed measurable but incomplete overlap between β-catenin chromobody and E-cadherin-positive regions. The mean thresholded Manders values were tM1 = 0.208 ± 0.096 and tM2 = 0.291 ± 0.107 (Fig. 3F). Together, these data support the presence of a localized E-cadherin-associated β-catenin pool in the mature pronephric epithelium, while also confirming that β-catenin remains distributed across additional compartments, including nuclei and cytosolic structures.

### 3.5 Differential nuclear localization of the β-catenin chromobody in proximal and distal mature tubules

To determine whether the β-catenin chromobody is sufficiently sensitive to detect biologically relevant differences in nuclear β-catenin localization within the kidney, we compared nuclear β-catenin chromobody fluorescence between the proximal and distal tubules of the mature pronephric kidney. Previous studies in vertebrate kidneys have demonstrated that canonical Wnt/β-catenin signaling is not uniform throughout the nephron and plays distinct roles in nephron patterning, segment specification, and maintenance of proximal tubule identity (21, 23, 49, 50). Therefore, differences in nuclear β-catenin localization between nephron segments would provide an independent assessment of the ability of the chromobody to report biologically meaningful changes in β-catenin distribution.

By stage NF40, the *Xenopus* pronephric kidney has formed well-organized epithelial tubules. At this stage, β-catenin chromobody signal was detected in the nuclei of both proximal and distal tubular cells (Fig 4A-B). Proximal tubules were identified using the 3G8 antibody, which labels the proximal tubular lumen and clearly delineates the proximal nephron segment (Fig. 4C). Distal tubules were identified using the 4A6 antibody, which specifically labels distal epithelial tubules (Fig. 4D). Although both antibodies were generated in the same species and therefore could not be simultaneously detected using different secondary antibodies, the distinctive lumen-restricted staining pattern of 3G8 enabled reliable identification and discrimination of the proximal and distal nephron segments.

**Figure 4.**
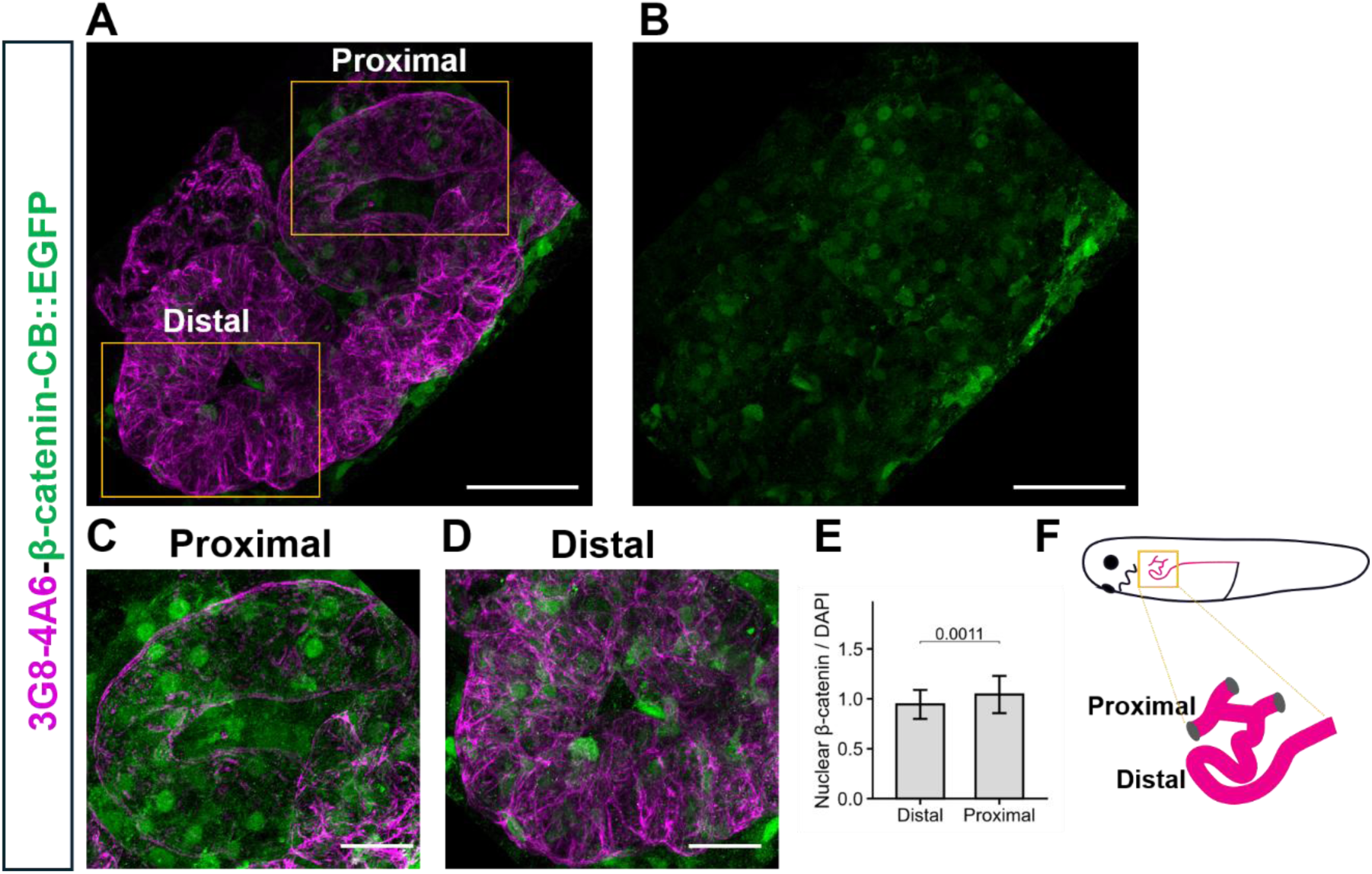
β-catenin chromobody localization in proximal and distal tubules of the mature pronephric kidney. At NF40, the pronephric kidney has formed organized tubule structures. β-catenin chromobody signal was detected in both proximal and distal tubules. Proximal tubules were labeled with 3G8, which stains the lumen and clearly delimits the proximal tubule region. Distal tubules were labeled with 4A6, showing epithelial distal tubule structures. Nuclear β-catenin chromobody intensity was measured within DAPI-defined nuclear ROIs located in proximal and distal tubule regions and normalized to DAPI intensity for each nucleus. Quantification showed higher nuclear β-catenin/DAPI levels in proximal nuclei compared with distal nuclei. Proximal and distal values were compared using a Wilcoxon rank-sum test, and the p-value is shown on the graph. Values are shown as mean ± SD. A diagram in F shows the localization of the mature nephritic kidney in a *Xenopus* tadpole. The magnification shows the proximal and distal structures evaluated in this experiment. Scale bar: 50 µm in panel A and B and 25 µm in panel C and D

Quantitative analysis of nuclear β-catenin chromobody fluorescence revealed significantly higher nuclear β-catenin levels in proximal tubule cells than in distal tubule cells (Fig. 4E). These findings indicate that, although nuclear β-catenin persists in the mature pronephric kidney, its distribution is not uniform along the nephron. Instead, β-catenin exhibits segment-specific nuclear enrichment, with significantly greater accumulation in proximal tubule nuclei than in distal tubule nuclei. These results are consistent with previous studies demonstrating spatial differences in Wnt/β-catenin activity during nephron development and support the β-catenin chromobody’s ability to detect physiologically relevant differences in β-catenin localization *in vivo*.

## 4. Discussion

### 4.1 The β-catenin chromobody suggests dynamic subcellular localization during Xenopus kidney development

We characterized the subcellular localization of β-catenin throughout *Xenopus* pronephric kidney development using an accelerated-turnover β-catenin chromobody. Although fluorescent chromobodies have previously been used to visualize endogenous proteins in living cells and other vertebrates, such as zebrafish (31, 37), their application to vertebrate kidney development had not been explored. Our study shows that the chromobody detects β-catenin in multiple cellular compartments, including nuclei, cytoplasm, and epithelial junctions, and suggests dynamic changes in localization during nephrogenesis. We first validated the chromobody in embryonic tissues with well-established β-catenin localization patterns. During gastrulation, strong nuclear fluorescence was detected in the dorsal organizer, consistent with the central role of canonical Wnt/β-catenin signaling in dorsal axis specification. In addition, the chromobody detected β-catenin enrichment within the embryonic epidermis, particularly in multiciliated cells, where β-catenin contributes to epithelial differentiation and ciliogenesis (38). These observations demonstrate that the chromobody shows β-catenin localization in tissues where its distribution has been characterized, supporting its use for subsequent analyses of kidney development.

### 4.2 Nuclear β-catenin predominates during early nephrogenesis

One of the principal findings of this study is that endogenous β-catenin undergoes dynamic developmental partitioning during nephrogenesis. Although β-catenin became progressively enriched at epithelial junctions as nephron epithelialization proceeded, nuclear β-catenin remained a substantial component of the total β-catenin pool throughout development. During late neurula and early tailbud stages (NF22 and NF26, respectively), β-catenin was enriched within the nuclei of Lhx1-positive nephron progenitor cells. Quantitative analyses demonstrated that nuclear β-catenin remained a substantial component of the total β-catenin pool throughout nephrogenesis, whereas epithelial-associated β-catenin became progressively more prominent during tubule maturation. These observations indicate that epithelial maturation is accompanied by coordinated expansion of multiple functional β-catenin pools rather than a simple redistribution from nuclear to junctional compartments.

These developmental changes indicate that increasing junctional recruitment appears to occur without obvious depletion of the nuclear β-catenin pool. This finding is consistent with the multifunctional roles of β-catenin during nephrogenesis, in which transcriptional and adhesive functions may coexist as the epithelial architecture matures. Canonical Wnt/β-catenin signaling promotes nephron induction, maintenance of nephron progenitors, and early epithelial differentiation in both amphibian and mammalian kidneys (21, 23, 50). Nuclear accumulation of β-catenin in Lhx1-positive progenitors is therefore consistent with transcriptional responses required for nephron specification and early epithelialization as a requirement for tubule formation. It is important to mention that although nuclear localization is frequently interpreted as evidence of canonical Wnt signaling, nuclear β-catenin alone does not necessarily indicate a positive transcriptional status. β-catenin continuously localizes between the cytoplasm and nuclear compartments, and only a subset of nuclear β-catenin participates in transcriptional regulation as a cofactor of regulators with TCF/LEF proteins. Moreover, the chromobody reports protein localization rather than transcriptional activation itself. Future studies combining the chromobody with transcriptional Wnt reporters (for example, the Wnt reporter frog, Xla.Tg(pbin7LEF-GFP)) will be valuable for distinguishing these two processes.

### 4.3 β-catenin progressively accumulates at epithelial junctions during tubule maturation

Importantly, increasing epithelial-associated localization did not occur at the expense of the nuclear pool. Instead, our quantitative analyses suggest that epithelial differentiation is accompanied by coordinated expansion of the adhesive β-catenin pool while maintaining substantial nuclear β-catenin enrichment. Our data showed that as epithelial differentiation progressed, β-catenin remained enriched in nuclei while progressively accumulating at epithelial cell borders. However, by stage NF40, fluorescence was detected along membrane structures throughout the mature pronephric tubules, while nuclear β-catenin remained detectable but represented a smaller proportion of the total signal. This redistribution is consistent with the dual functions of β-catenin, which means that, in addition to acting as a transcriptional co-activator, β-catenin is a structural component of adherens junctions, where it links classical cadherins to the actin cytoskeleton and contributes to epithelial integrity (51–53). As nephron epithelialization proceeds, the recruitment of β-catenin to adherens junctions is expected to increase as stable epithelial architecture becomes established. Colocalization analysis (54, 55) further supports this interpretation and demonstrates a moderate positive Pearson correlation between β-catenin chromobody and E-cadherin. Thresholded Manders coefficients indicated that only a subset of β-catenin overlapped with E-cadherin-positive membranes, consistent with the presence of multiple intracellular β-catenin pools. Rather than being restricted to epithelial junctions, β-catenin remained detectable in the nuclei and cytoplasm of mature tubules, reflecting its diverse subcellular functions.

### 4.4 Nuclear β-catenin differs between mature nephron segments

Although junctional localization became increasingly prominent during epithelial maturation, nuclear β-catenin remained abundant and exhibited segment-specific differences within the mature nephron. Quantification of nuclear fluorescence demonstrated significantly greater nuclear β-catenin in proximal tubule cells than in distal tubule cells. Previous studies have established that canonical Wnt signaling is spatially regulated during nephron patterning and differentiation (21, 23, 49, 50). Our findings are consistent with continued spatial regulation of β-catenin partitioning after nephron morphogenesis, with significantly greater nuclear enrichment in proximal than distal tubules. These observations suggest that distinct nephron segments may maintain different levels of canonical Wnt signaling or β-catenin turnover after epithelial differentiation. Whether these differences reflect persistent canonical Wnt signaling or distinct rates of β-catenin turnover across development, and whether they involve segment-specific regulation of nuclear import and export, remains to be determined. These data indicate that β-catenin localization remains regionally regulated after nephron morphogenesis is largely complete.

### 4.5 Limitations and future applications

Although the chromobody provides a powerful approach for visualizing endogenous β-catenin localization, localization alone does not establish functional activity or distinguish transcriptionally active from inactive nuclear β-catenin. Future studies combining the chromobody with functional perturbations, canonical Wnt reporters, or transcriptional readouts will help determine how developmental changes in β-catenin partitioning relate to signaling activity and nephron morphogenesis.

The temporal resolution of our live imaging was also limited. Although the chromobody enabled visualization of β-catenin localization throughout embryogenesis, live imaging of the pronephric kidney presented several technical challenges. The abundance of yolk platelets and melanocytes in later-stage embryos reduced optical penetration. Future improvements in imaging approaches, including albino *Xenopus* lines and light-sheet microscopy, should further improve visualization of β-catenin dynamics during nephrogenesis. Mosaic chromobody expression resulting from blastomere microinjection likely contributed to variability in fluorescence intensity among kidney cells. Stable transgenic reporter lines should further improve quantitative analyses of β-catenin dynamics. Despite these limitations, the β-catenin chromobody provides an unprecedented opportunity to visualize endogenous β-catenin dynamics throughout vertebrate nephrogenesis *in vivo*.

Together, these findings support a developmental model in which epithelial maturation is accompanied by the coordinated partitioning of β-catenin among nuclear, cytoplasmic, and adhesive compartments, rather than by a simple exchange between transcriptional and structural pools. By revealing how these intracellular β-catenin populations change during nephrogenesis, this work provides a framework for understanding how canonical Wnt signaling is integrated with epithelial morphogenesis.

## Acknowledgements

We thank the members of the laboratories of Dr. Jae-il Park and Dr. Malgorzata Kloc, as well as Dr. Jun Wang, Dr. Kendra Carmon, and Dr. George Eisenhoffer, for their constructive feedback, encouragement, helpful discussions, and scientific advice throughout this project. We are especially grateful to Dr. Yoshihiro Komatsu and Dr. Hiro Yamaguchi for providing access to equipment used in this study. We thank the animal care technicians and veterinary staff, including J.C. Whitney, Amy Campbell, Jamie Cochran, and Thomas Gomez, for their excellent care of the *Xenopus* colony. We also acknowledge the National *Xenopus* Resource for providing the transgenic *Xenopus* lines used in this work. We are particularly grateful to Dr. Peter Walentek for his invaluable guidance and insightful discussions on Wnt signaling and β-catenin biology, which greatly contributed to the interpretation of this study.

## Author contributions

RKM conceptualized this project. UR provided the chromobody and advice to optimize its protein turnover. Chromobody experiments were performed by AR, ACM, and BLW. Figures assembly was carried out by AR, ACM, and BLW. Data analysis was performed by AR. The manuscript was written by AR. Project administration, as well as data and manuscript evaluation, were carried out by RKM.

## References

1. Schmidt-Ott KM, Lan D, Hirsh BJ, Barasch J. Dissecting stages of mesenchymal-to-epithelial conversion during kidney development. Nephron Physiol. 2006;104(1):p56–60.

2. Zhang P, Cai Y, Soofi A, Dressler GR. Activation of Wnt11 by Transforming Growth Factor-beta Drives Mesenchymal Gene Expression through Non-canonical Wnt Protein Signaling in Renal Epithelial Cells. J Biol Chem. 2012;287(25):21290–302.

3. Wessely O, Tran U. Xenopus pronephros development--past, present, and future. Pediatr Nephrol. 2011;26(9):1545–51.

4. Corkins ME, Achieng M, DeLay BD, Krneta-Stankic V, Cain MP, Walker BL, et al. A comparative study of cellular diversity between the Xenopus pronephric and mouse metanephric nephron. Kidney Int. 2023;103(1):77–86.

5. Corkins ME, Romero-Mora A, Achieng MA, Lindstrom NO, Miller RK. Comparative analysis of Xenopus mesonephric transcriptomics: Conservation of the developmental lineage of nephron stages. bioRxiv. 2025.

6. Raciti D, Reggiani L, Geffers L, Jiang Q, Bacchion F, Subrizi AE, et al. Organization of the pronephric kidney revealed by large-scale gene expression mapping. Genome Biol. 2008;9(5):R84.

7. Zhou X, Vize PD. Proximo-distal specialization of epithelial transport processes within the Xenopus pronephric kidney tubules. Dev Biol. 2004;271(2):322–38.

8. Corkins ME, Krneta-Stankic V, Kloc M, Miller RK. Aquatic models of human ciliary diseases. Genesis. 2021;59(1-2):e23410.

9. Blackburn ATM, Miller RK. Modeling congenital kidney diseases in Xenopus laevis. Dis Model Mech. 2019;12(4).

10. Corkins ME, Hanania HL, Krneta-Stankic V, DeLay BD, Pearl EJ, Lee M, et al. Transgenic Xenopus laevis Line for In Vivo Labeling of Nephrons within the Kidney. Genes (Basel). 2018;9(4).

11. Krneta-Stankic V, Corkins ME, Paulucci-Holthauzen A, Kloc M, Gladden AB, Miller RK. The Wnt/PCP formin Daam1 drives cell-cell adhesion during nephron development. Cell Rep. 2021;36(1):109340.

12. Maher MT, Flozak AS, Stocker AM, Chenn A, Gottardi CJ. Activity of the beta-catenin phosphodestruction complex at cell-cell contacts is enhanced by cadherin-based adhesion. J Cell Biol. 2009;186(2):219–28.

13. Mo R, Chew TL, Maher MT, Bellipanni G, Weinberg ES, Gottardi CJ. The terminal region of beta-catenin promotes stability by shielding the Armadillo repeats from the axin-scaffold destruction complex. J Biol Chem. 2009;284(41):28222–31.

14. Reinacher-Schick A, Gumbiner BM. Apical membrane localization of the adenomatous polyposis coli tumor suppressor protein and subcellular distribution of the beta-catenin destruction complex in polarized epithelial cells. J Cell Biol. 2001;152(3):491–502.

15. Roberts DM, Pronobis MI, Poulton JS, Waldmann JD, Stephenson EM, Hanna S, Peifer M. Deconstructing the sscatenin destruction complex: mechanistic roles for the tumor suppressor APC in regulating Wnt signaling. Mol Biol Cell. 2011;22(11):1845–63.

16. Piao S, Lee SH, Kim H, Yum S, Stamos JL, Xu Y, et al. Direct inhibition of GSK3beta by the phosphorylated cytoplasmic domain of LRP6 in Wnt/beta-catenin signaling. PLoS One. 2008;3(12):e4046.

17. Nelson WJ, Nusse R. Convergence of Wnt, beta-catenin, and cadherin pathways. Science. 2004;303(5663):1483–7.

18. Nelson WJ, Veshnock PJ. Ankyrin binding to (Na+, K+)ATPase and implications for the organization of membrane domains in polarized cells. Nature. 1987;328:533–6.

19. Valenta T, Lukas J, Doubravska L, Fafilek B, Korinek V. HIC1 attenuates Wnt signaling by recruitment of TCF-4 and beta-catenin to the nuclear bodies. EMBO J. 2006;25(11):2326–37.

20. Park JS, Ma W, O’Brien LL, Chung E, Guo JJ, Cheng JG, et al. Six2 and Wnt regulate self-renewal and commitment of nephron progenitors through shared gene regulatory networks. Dev Cell. 2012;23(3):637–51.

21. Park JS, Valerius MT, McMahon AP. Wnt/beta-catenin signaling regulates nephron induction during mouse kidney development. Development. 2007;134(13):2533–9.

22. Karner CM, Chirumamilla R, Aoki S, Igarashi P, Wallingford JB, Carroll TJ. Wnt9b signaling regulates planar cell polarity and kidney tubule morphogenesis. Nat Genet. 2009;41(7):793–9.

23. Karner CM, Das A, Ma Z, Self M, Chen C, Lum L, et al. Canonical Wnt9b signaling balances progenitor cell expansion and differentiation during kidney development. Development. 2011;138(7):1247–57.

24. Saulnier DM, Ghanbari H, Brandli AW. Essential function of Wnt-4 for tubulogenesis in the Xenopus pronephric kidney. Dev Biol. 2002;248(1):13–28.

25. Lyons JP, Miller RK, Zhou X, Weidinger G, Deroo T, Denayer T, et al. Requirement of Wnt/beta-catenin signaling in pronephric kidney development. Mech Dev. 2009;126(3-4):142–59.

26. Lyons JP, Mueller UW, Ji H, Everett C, Fang X, Hsieh JC, et al. Wnt-4 activates the canonical beta-catenin-mediated Wnt pathway and binds Frizzled-6 CRD: functional implications of Wnt/beta-catenin activity in kidney epithelial cells. Exp Cell Res. 2004;298(2):369–87.

27. Murugan S, Shan J, Kuhl SJ, Tata A, Pietila I, Kuhl M, Vainio SJ. WT1 and Sox11 regulate synergistically the promoter of the Wnt4 gene that encodes a critical signal for nephrogenesis. Exp Cell Res. 2012;318(10):1134–45.

28. Garriock RJ, Warkman AS, Meadows SM, D’Agostino S, Krieg PA. Census of vertebrate Wnt genes: isolation and developmental expression of Xenopus Wnt2, Wnt3, Wnt9a, Wnt9b, Wnt10a, and Wnt16. Dev Dyn. 2007;236(5):1249–58.

29. Majumdar A, Vainio S, Kispert A, McMahon J, McMahon AP. Wnt11 and Ret/Gdnf pathways cooperate in regulating ureteric branching during metanephric kidney development. Development. 2003;130(14):3175–85.

30. O’Brien LL, Combes AN, Short KM, Lindstrom NO, Whitney PH, Cullen-McEwen LA, et al. Wnt11 directs nephron progenitor polarity and motile behavior ultimately determining nephron endowment. Elife. 2018;7.

31. Rothbauer U, Zolghadr K, Tillib S, Nowak D, Schermelleh L, Gahl A, et al. Targeting and tracing antigens in live cells with fluorescent nanobodies. Nat Methods. 2006;3(11):887–9.

32. Traenkle B, Emele F, Anton R, Poetz O, Haeussler RS, Maier J, et al. Monitoring interactions and dynamics of endogenous beta-catenin with intracellular nanobodies in living cells. Mol Cell Proteomics. 2015;14(3):707–23.

33. Nieuwkoop PD, Faber J. Normal table of Xenopus laevis (Daudin) : a systematical and chronological survey of the development from the fertilized egg till the end of metamorphosis. New York: Garland Pub.; 1994. 252 p., 10 leaves of plates p.

34. Gauvrit S, Zhao S, Rothbauer U, Stainier DYR. A beta-catenin chromobody-based probe highlights endothelial maturation during vascular morphogenesis in vivo. Development. 2024;151(11).

35. Traenkle B, Rothbauer U. Under the Microscope: Single-Domain Antibodies for Live-Cell Imaging and Super-Resolution Microscopy. Front Immunol. 2017;8:1030.

36. Venegas-Ferrin M, Sudou N, Taira M, del Pino EM. Comparison of Lim1 expression in embryos of frogs with different modes of reproduction. Int J Dev Biol. 2010;54(1):195–202.

37. Panza P, Maier J, Schmees C, Rothbauer U, Sollner C. Live imaging of endogenous protein dynamics in zebrafish using chromobodies. Development. 2015;142(10):1879–84.

38. Haas M, Gomez Vazquez JL, Sun DI, Tran HT, Brislinger M, Tasca A, et al. DeltaN-Tp63 Mediates Wnt/beta-Catenin-Induced Inhibition of Differentiation in Basal Stem Cells of Mucociliary Epithelia. Cell Rep. 2019;28(13):3338–52 e6.

39. Keller BM, Maier J, Secker KA, Egetemaier SM, Parfyonova Y, Rothbauer U, Traenkle B. Chromobodies to Quantify Changes of Endogenous Protein Concentration in Living Cells. Mol Cell Proteomics. 2018;17(12):2518–33.

40. Winklbauer R, Nagel M, Selchow A, Wacker S. Mesoderm migration in the Xenopus gastrula. Int J Dev Biol. 1996;40(1):305–11.

41. Winklbauer R, Selchow A. Motile behavior and protrusive activity of migratory mesoderm cells from the Xenopus gastrula. Dev Biol. 1992;150(2):335–51.

42. Winklbauer R, Selchow A, Nagel M, Angres B. Cell interaction and its role in mesoderm cell migration during Xenopus gastrulation. Dev Dyn. 1992;195(4):290–302.

43. Myers DC, Sepich DS, Solnica-Krezel L. Convergence and extension in vertebrate gastrulae: cell movements according to or in search of identity? Trends Genet. 2002;18(9):447–55.

44. Skoglund P, Rolo A, Chen X, Gumbiner BM, Keller R. Convergence and extension at gastrulation require a myosin IIB-dependent cortical actin network. Development. 2008;135(14):2435–44.

45. Keller R, Davidson L, Edlund A, Elul T, Ezin M, Shook D, Skoglund P. Mechanisms of convergence and extension by cell intercalation. Philos Trans R Soc Lond B Biol Sci. 2000;355(1399):897–922.

46. Zoltewicz JS, Gerhart JC. The Spemann organizer of Xenopus is patterned along its anteroposterior axis at the earliest gastrula stage. Dev Biol. 1997;192(2):482–91.

47. Vonica A, Gumbiner BM. The Xenopus Nieuwkoop center and Spemann-Mangold organizer share molecular components and a requirement for maternal Wnt activity. Dev Biol. 2007;312(1):90–102.

48. Ochi H, Tamai T, Nagano H, Kawaguchi A, Sudou N, Ogino H. Evolution of a tissue-specific silencer underlies divergence in the expression of pax2 and pax8 paralogues. Nat Commun. 2012;3:848.

49. Lindstrom NO, Lawrence ML, Burn SF, Johansson JA, Bakker ER, Ridgway RA, et al. Integrated beta-catenin, BMP, PTEN, and Notch signalling patterns the nephron. Elife. 2015;3:e04000.

50. Carroll TJ, Park JS, Hayashi S, Majumdar A, McMahon AP. Wnt9b plays a central role in the regulation of mesenchymal to epithelial transitions underlying organogenesis of the mammalian urogenital system. Dev Cell. 2005;9(2):283–92.

51. Huber AH, Stewart DB, Laurents DV, Nelson WJ, Weis WI. The cadherin cytoplasmic domain is unstructured in the absence of beta-catenin. A possible mechanism for regulating cadherin turnover. J Biol Chem. 2001;276(15):12301–9.

52. Kemler R. From cadherins to catenins: Cytoplasmic protein interactions and regulation of cell adhesion. Trends Genet. 1993;9:317–21.

53. Drees F, Pokutta S, Yamada S, Nelson WJ, Weis WI. Alpha-catenin is a molecular switch that binds E-cadherin-beta-catenin and regulates actin-filament assembly. Cell. 2005;123(5):903–15.

54. Costes SV, Daelemans D, Cho EH, Dobbin Z, Pavlakis G, Lockett S. Automatic and quantitative measurement of protein-protein colocalization in live cells. Biophys J. 2004;86(6):3993–4003.

55. Aaron JS, Taylor AB, Chew TL. Image co-localization - co-occurrence versus correlation. J Cell Sci. 2018;131(3).

