## Supplementary material for "Developmental shift in β-catenin localization between nuclear and junctional pools during vertebrate nephron development": S1 AND S2

### Supplemental files

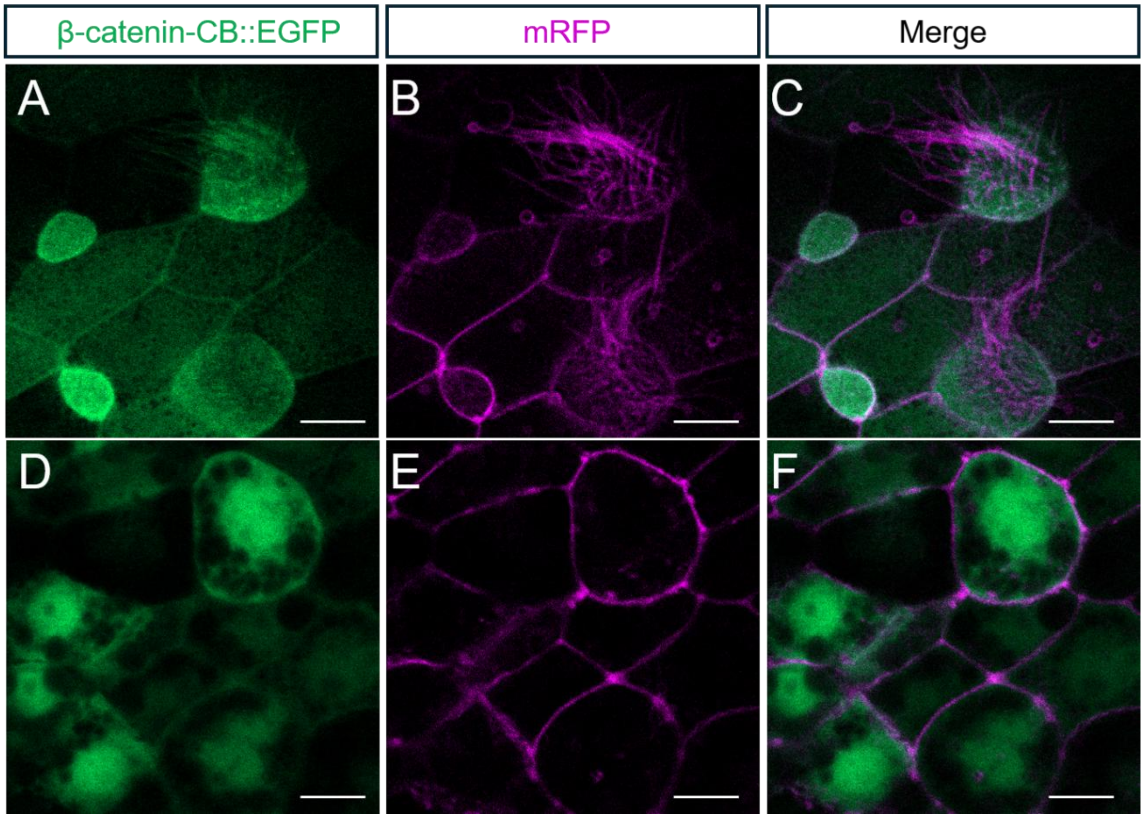


**Figure S1. β-catenin chromobody localization in the epidermis of *Xenopus* embryos.**
Live imaging of *Xenopus* embryos injected with β-catenin-CB::EGFP mRNA and mRFP tracer. β-catenin-CB::EGFP is shown in green, and the mRFP tracer is shown in magenta. Panels A, B, and C at the top show the early tailbud epidermis, depicting multiciliated cells alongside the β-catenin chromobody. The merge shows the relationship between β-catenin chromobody signal and tracer-labeled cells. Panels D–F show deeper optical sections with β-catenin signal in multiciliated cells and cytoplasmic β-catenin signal in neighboring cells. Scale bar: 10 μm.

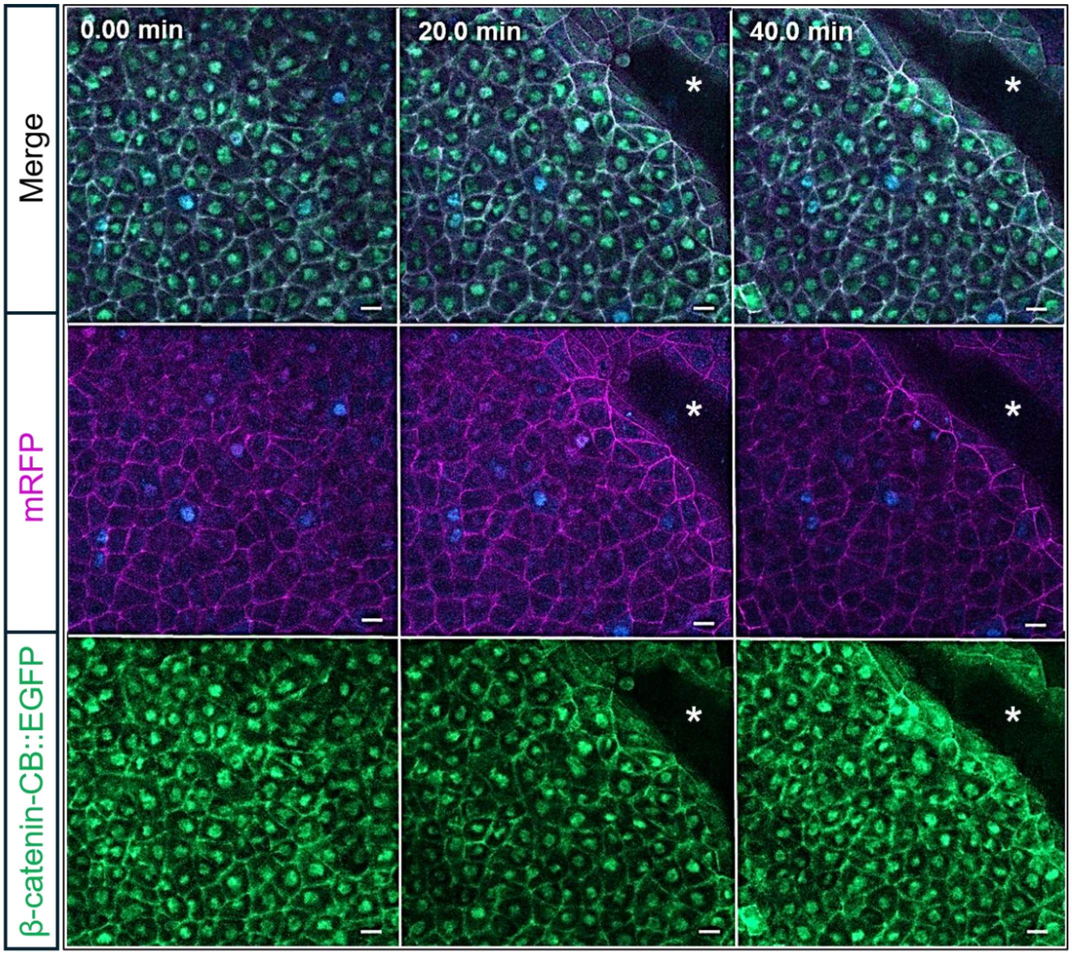


**Figure S2. β-catenin chromobody localization in *X. laevis* early gastrula embryos in the dorsal lip during blastopore closure.**

Live imaging of early gastrula-stage *X. laevis* embryos injected with β-catenin-CB::EGFP mRNA and a membrane tracer RFP. β-catenin-CB::EGFP labeling endogenous β-catenin is shown in green, the membrane tracer is shown in magenta, and nuclei are labeled with Hoechst in blue. The β-catenin chromobody signal is detected near the dorsal blastopore lip (asterisk), including nuclear and cell border regions, consistent with β-catenin localization during dorsal organizer formation. Scale bar: 10 µm.
